# Sustained Activation of a Defensive Behavior-Related Circuit Elicits a distinct Homeostatic Sleep Response

**DOI:** 10.64898/2026.10.09.758035

**Authors:** Chun Yang, Victoria Lombardi, David S. Uygun, Ritchie E. Brown, James T. McKenna, Radhika Basheer

## Abstract

Animals and humans often forgo sleep in response to other physiological needs. Prolonged wakefulness leads to a compensatory homeostatic sleep response involving enhanced amounts of sleep and electroencephalographic delta (0.5-4 Hz) power, yet how the type of preceding wakefulness and brain activity affects this response is poorly understood. Here we compared the pattern of brain activity and the homeostatic sleep response following sustained wakefulness induced by either gentle handling (GH), a widely used form of sleep deprivation in rodents; or optogenetic stimulation of neurons involved in avoidance/defensive responses, basal forebrain (BF) vesicular glutamatergic transporter type 2 (vGluT2) neurons. Local infusion of ionotropic glutamate receptor antagonists partially attenuated the wake-inducing effect of vGluT2-stimulation, but not that of GH, suggesting mediation by other non-glutamatergic BF subtypes. These antagonists also tended to produce more NREM sleep and a shorter NREM latency in the recovery period following vGluT2-stimulation. Both manipulations increased high theta (7-9 Hz) during wakefulness, an effect partially blocked by local infusion of the glutamate antagonist cocktail. Enhanced NREM sleep time and NREM delta power in the recovery period were more pronounced following GH than following vGluT2 stimulation. Whole-brain cFos immunohistochemistry revealed stronger neuronal activity in brain regions involved in stress and avoidance/defensive behaviors with vGluT2 stimulation, compared to GH. Our results demonstrate that activation of a brain system involved in defensive behavior (BF vGluT2) elicits a distinct and diminished homeostatic sleep response compared to GH, potentially an evolutionary sleep adaptation which would maintain vigilance in the face of an ongoing threat.

## Introduction

Sleep is present throughout the animal kingdom and considered essential for health and survival ^1,2^. Nonetheless, animals and humans often adaptively curtail sleep in the presence of other important physiological drives such as the presence of rewarding stimuli or threats to survival ^3–5^. The resulting prolonged wakefulness normally elicits a homeostatic sleep response (HSR) characterized by increased cortical theta (4-8 Hz) activity during the period of sleep deprivation followed by sleep that is typically longer in duration and deeper in intensity, with increased electroencephalogram delta (0.5–4 Hz) power during non-rapid-eye-movement (NREM) sleep ^6–10^. However, increasing evidence suggests that the expression of the HSR depends on the type of preceding wakefulness and the corresponding pattern of brain activity. For instance, in rodents, wakefulness produced by wheel running or engagement in a well-trained touchscreen task motivated by food reward, produces a reduced HSR compared to wakefulness produced by exposure to novel objects ^11^. Furthermore, activation of different arousal pathways elicits different homeostatic sleep responses ^12–18^. Thus, much remains to be studied regarding the inter-relationship between different patterns of brain activity and the subsequent HSR.

Investigation of the HSR following activation of brain systems involved in responses to threat is of translational importance given the relationship between these systems and clinically observed sleep disorders such as insomnia and PTSD ^19–21^. Thus, in this study we used EEG analysis and whole-brain mapping of the immediate early gene product, cFos, to compare the pattern of brain activity and the HSR between prolonged wakefulness produced by optogenetic activation of a brain system involved in avoidance and other defensive behaviors related to anxiety and depression ^22–26^, basal forebrain (BF) vesicular glutamatergic transporter type 2 (vGluT2) neurons, and a commonly used means of sleep deprivation in rodent studies, gentle handling (GH). Furthermore, using reverse microdialysis of glutamate receptor antagonists, we dissected the role of BF glutamatergic receptors in different aspects of the HSR. Both GH and vGluT2-stimulation kept the animals awake to similar extent. Although BF vGluT2 stimulation promoted sustained wakefulness, it resulted in a weaker HSR and preferentially activated a network of avoidance/defensive behavior-related brain regions, compared to GH. The homeostatic regulation of sleep therefore depends not only on the preceding time awake, but also which specific neural circuits are engaged to promote wakefulness. In particular, the distinct HSR following prolonged wakefulness produced by activation of BF vGluT2 neurons, a brain system involved in avoidance behaviors, potentially serves as an evolutionary adaptation which maintains enhanced vigilance in the face of a perceived or ongoing threat.

## Results

### Opto-stimulation of BF vGluT2 neurons (vGluT2-stim) or gentle handling (GH) produce sustained wakefulness (**Fig. 1**)

vGluT2-Cre mice (N=10) and C57Bl6 mice (N=7) were implanted with EEG/EMG electrodes. vGluT2-Cre mice were injected with AAV-DIO-ChR2-eYFP into BF and used for experiments after 4 weeks. A random and counter-balanced sequence of different testing conditions (GH or vGluT2 stim, drug infusion, ACSF infusion or baseline) was applied in the experimental design to avoid order effects. Both genotypes were recorded for 24 h to determine baseline (BL) sleep-wakefulness. vGluT2 mice received opto-stimulation of BF vGluT2 neurons (Stim; 5s ON: 55s OFF) from 10AM to 2PM (**Fig 1A**). C57Bl6 mice (N=7) underwent sleep deprivation by GH from 10AM -2PM (ZT3-7). The BL recordings of both genotypes showed no differences. Opto-stimulations of vGluT2 neurons led to immediate arousal responses, as noted by low voltage, fast-frequency EEG and increased muscle activity at the onset of optical stimulation (**Fig 1B).** Compared to the baseline day (BL wake%: 30.85±1.67%, N=10), GH and vGluT2-stim significantly increased wakefulness (wake%: GH: 99.25±0.45%, N=7, p<0.001 compared to BL; vGluT2-stim: 93.31±1.23%, N=10, p<0.05 compared to BL), with no significant difference between GH and vGluT2-stim (p>0.05; ANOVA with Bonferroni’s comparison test). Thus, increases in the time spent in wakefulness during the 4 h of BF vGlut2 stimulation was comparable to those noted for GH. Nonetheless, there was a small but gradual decrease in hourly wakefulness in the vGluT2-stim condition (hour1 wake%: 99.34 ± 0.32%, hour 2 wake%:96.40 ± 0.83%; hour 3 wake%: 91.42 ± 2.03%, p<0.01 compared to hour 1; hour 4 wake%: 86.09 ± 2.69%, p<0.001 compared to hour 1, p<0.01 compared to hour 2 or 3. Repeated measures ANOVA with Bonferroni’s comparison test) (**Fig 1C**), suggesting an increased pressure to sleep or a diminished ability of BF vGluT2 neurons to promote wakefulness. The power spectral analysis of the EEG for the 4h revealed distinct frequency-specific differences between vGluT2-stim and GH conditions. When compared to BL, vGluT2-stim elicited a significant increase in the 0.5-3Hz (delta range, p<0.05) and 7-13Hz (high theta and sigma range, p<0.05), whereas during GH, an increase in 1.5Hz and 7-15Hz frequencies (p<0.05) in the wake EEG was observed (**Fig 1D**). Post-hoc histology of AAV-ChR2 injected vGluT2-Cre mice confirmed the location of the cannula tip to be within the transduced glutamatergic population of the BF **(Fig 1E**).

**Figure 1:**
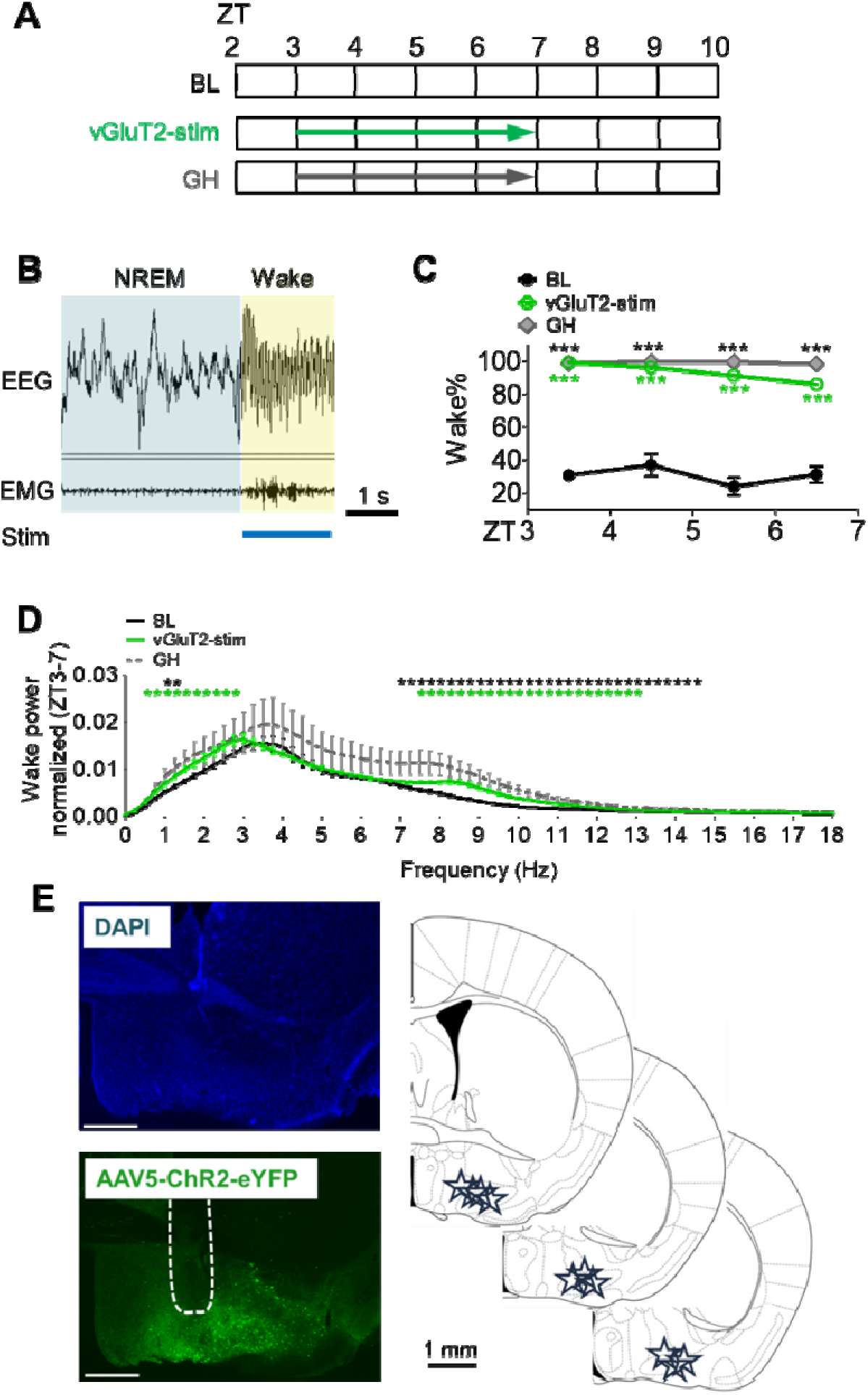
Optostimulation of BF vGluT2^+^ neurons (vGluT2-stim) induced rapid and prolonged wakefulness. **A**: Experimental protocol for three different recording paradigms: baseline day (BL: no stimulation), optostimulation day (vGluT2-stim: blue light of 20mW/cm^2^ was applied to the right BF at 20Hz with 10 msec pulses and 5 s of light per minute from ZT3-7), and acute sleep deprivation by gentle handling (GH: from ZT3-7). ZT: zeitgeber time. Light was on at ZT0 (7AM). **B**. A representative sleep recording trace showing an immediate transition from NREM to wakefulness with the optostimulation of BF vGluT2^+^ neurons. EEG: electroencephalogram. EMG: electromyogram. Stim: vGluT2-stim with blue light (indicated by blue bar). NREM: non-rapid eye movement. **C**: Hourly percentage time spent in wakefulness from ZT3-7 during spontaneous sleep (baseline, BL, N=10), BF vGluT2^+^ -stim (N=10) and GH (N=7). There was no significant difference between vGluT2-stim and GH. ***: p<0.001, ANOVA with Bonferroni’s comparison test. Green asterisk: vGluT2-stim vs. BL. Black asterisk: GH vs. BL. **D:** Wake spectral analysis showed BF vGluT2-stim led to a significant increase in 0.5-3Hz (delta range) and 7-13Hz (high theta and sigma range), whereas during GH, an increase in 1-1.5Hz and 7-15Hz frequencies in EEG was observed when compared to BL . *: p<0.05, t-test. Green asterisk: vGluT2-stim vs BL. Black asterisk: GH vs BL. **E**: Images and sketches show the target locations of AAV5-ChR2-eYFP injection and expression and optical probe implantation within BF, (star, N=10) in vGluT2-cre mice. Dash line on the image shows the trace of microdialysis probe targeting BF. Scale bar: 500 µm.

### Prolonged wakefulness elicited by BF vGluT2-stim was partially mediated by activation of local ionotropic glutamate receptors in BF (**Fig. 2**)

BF receives robust glutamatergic input from multiple brain regions, including the lateral hypothalamus, the amygdala, parabrachial nucleus and the brain stem ^27–32^. The local glutamatergic neurons also extend collaterals and excite neighboring neurons ^33,34^. To evaluate the contribution of local BF glutamatergic signaling to arousal, we performed sleep deprivation by both vGluT2-stim and GH with and without the reverse microdialysis of a cocktail of ionotropic glutamate receptors’ (NMDA and AMPA) antagonists (DNQX and D-AP5) from ZT 3-10 (**Fig 2A**) using our previously described optodialysis probes ^35^. The antagonist cocktail treatment had no effect on spontaneous sleep-wakefulness (ZT3-7 wake%: 29.16 ± 4.62%. p=0.7343 compared to BL (ANOVA with Bonferroni’s comparison test). However, 4h of vGluT2-stim induced wakefulness was decreased to 76.66 ± 4.56% in the presence of an antagonist cocktail (p<0.001 compared to vGluT2-stim alone; p<0.0001 compared to drug) (**Fig. S1**). The decrease in hourly wakefulness was statistically significant starting from hour 2-4 compared to stimulation alone (Stim + antagonist: hour 1 wake%: 90.21±3.22%, hour 2 wake%: 79.98±5.01%, p<0.05 compared to stim hour 2; hour 3 wake%: 70.50±5.95%, p<0.05 compared to stim hour 3; hour 4 wake%: 65.94±5.34%, p<0.05 compared to stim hour 4 (**Fig 2 B1**). In contrast, the antagonists did not affect baseline wakefulness and interestingly had no significant effects on wakefulness induced by GH (**Fig 2 B2).**

**Figure 2:**
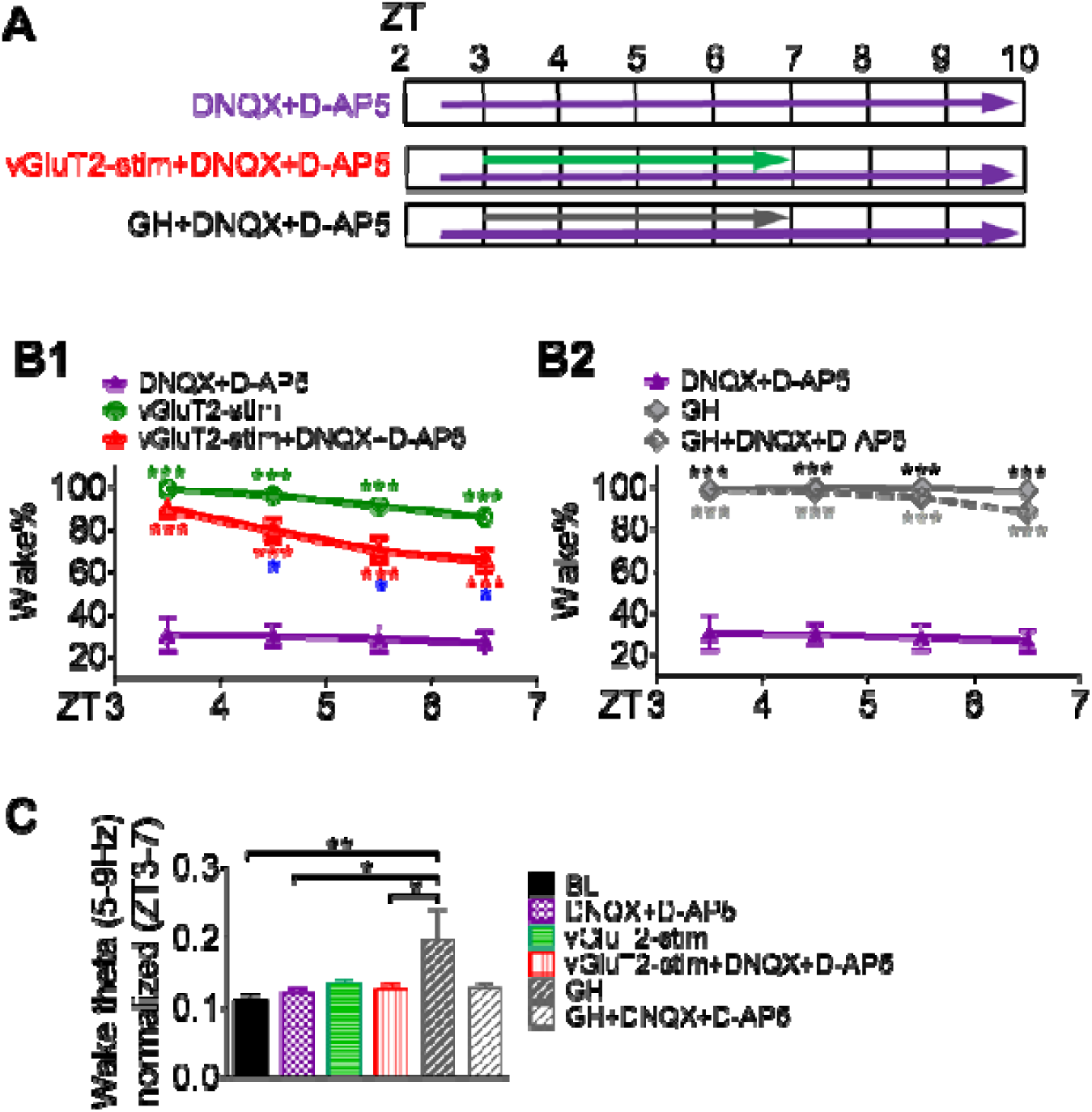
Prolonged wakefulness induced by optostimulation of BF vGluT2 neurons (vGluT2-stim) was partially mediated by activation of local ionotropic glutamate receptors in BF. **A**: Experimental protocol for three different conditions: drug infusion day (DNQX+D-AP5: A cocktail of ionotropic glutamatergic receptor antagonists 200 μM DNQX and 500 μM D-AP5 were infused into right BF via reverse microdialysis from ZT 2.5-10; N=10), vGluT2 stim+drug infusion day (vGluT2-stim+DNQX+D-AP5: vGluT2-stim from ZT3-7 with infusion of DNQX+D-AP5 from ZT2.5-10; N=10), and sleep deprivation by gentle handling (GH) + drug infusion day (GH+DNQX+D-AP5: SD from ZT3-7 with infusion of DNQX+D-AP5 from ZT2.5-10; N=7). ZT: Zeitgeber Time. **B**. Antagonism of ionotropic glutamate receptors in BF reduces the hourly percentage time spent in wakefulness induced by vGluT2-stim (**B1**) but not GH (**B2**). The drug alone had no effect on BL sleep (not shown). *: p<0.05, ***: p<0.001, ANOVA with Bonferroni’s comparison test. Green asterisk: vGlut2-stim vs. drug. Red asterisk: vGlut2-stim+drug vs. drug. Black asterisk: GH vs. drug. Grey asterisk: GH+drug vs. drug. Blue asterisk in B1: vGluT2-stim vs. vGluT2-stim+drug. **C:** Sleep deprivation by GH increased power of theta activity (5-9 Hz) in the wake state (ZT3-7), while vGluT2-stim induced mild but non-significant change. Local blockade of ionotropic glutamate receptors in BF eliminated the increase in theta activity induced by GH. *: p<0.05, **: p<0.01, ANOVA with Bonferroni’s comparison test.

The progressive increase in wake-theta (θ, 5-9Hz) activity is described as a marker of sleep homeostasis both in humans and in rodents ^36–39^. Therefore, we examined the changes in theta activity in our mice. There was no significant difference in the baseline wake-θ between vGluT2-stim and GH. Interestingly, during GH the wake θ power increased significantly (77.01% increase from BL, p<0.01, ANOVA with Bonferroni’s test), with no significant change with vGluT2-stim. The increased wake θ during GH was blocked by the local BF infusion of ionotropic glutamate receptor antagonists (**Fig 2C**). In rodents, low-frequency EEG theta activity (5–7 Hz) is associated with quiet waking and sleepiness ^38,39^, while high-frequency (7–9 Hz) is associated with active wakefulness and related cortical arousal ^38,40–44^. We observed that GH led to a trend-level increase in low theta (5-7 Hz), but not in the vGluT2-stim group (see Fig 1D), suggesting a weaker homeostatic theta response to vGluT2-Stim. However, in both conditions (GH and vGluT2-stim) high theta (7-9Hz) showed a significant increase from BL and the increases were blocked by the local administration of ionotropic glutamate receptor antagonists (**Fig. S2**).

### Sleep deprivation-induced by vGluT2-stim leads to a weaker homeostatic response compared to that of GH (**Figure 3**, S3)

Acute sleep deprivation elicits an HSR with increased NREM sleep duration and increased intensity in NREM EEG power in the delta range 0.5-4Hz (δ) ^6–9^. The baseline NREM δ spanning 0.5-4Hz was higher in the WT C57Bl6 mice used for GH when compared to vGluT2-Cre mice. Both genotypes showed significant increases in the recovery NREM δ spanning 0.5-4 Hz, but though statistically significant, it was less pronounced following BF vGluT2 stim compared to that observed after sleep deprivation with GH (Stim: 20.09% increase from BL, p<0.01 compared to BL; GH: 53.65% increase from BL, p<0.001 compared to BL. ANOVA with Bonferroni’s test). Recent reports suggest that higher frequency delta (between 2-4 Hz), defined as δ2 ^45^ is the homeostatic component which decays faster with ensuing sleep after prolonged waking ^45,46^. Both conditions showed significant increases in the delta activity in 2.5-3.5Hz range. Importantly, the normalized increase in δ2 was significantly higher with GH when compared to vGluT2-stim (t-test, p<0.0007). suggesting a stronger homeostatic response following GH. The antagonists increased the δ2 for both manipulations and were more pronounced and statistically significant (p<0.05, repeated-measures ANOVA with Bonferroni’s comparison) following vGluT2-stim compared to GH (**Fig 3 B, C**). It was intriguing that NREM sleep time in the 3 h (ZT7-10) that followed vGluT2 stimulation was not increased, suggesting that this aspect of the homeostatic sleep response was eliminated (Stim vs BL: p=0.9948. unpaired-t-test.). This observation was in contrast to the time-matched NREM sleep opportunity following 4 h of GH (GH NREM%: 66.58±0.92% vs BL NREM%: 60.56±1.99%. p=0.0305, unpaired t-test) (**Fig. S3A**). Instead, during the first hour of the recovery period, animals had significantly less sleep after 4h vGluT2-stim compared to BL (p=0.0112, t-test). This reduction of sleep was eliminated in the presence of ionotropic glutamate receptor antagonists (p=0.8459 compared to BL, t-test) (**Fig. S3B**). In addition, in the presence of glutamate receptor antagonists, at the cessation of opto-stimulation, the latency to NREM sleep decreased significantly (Latency after stim: 455.7 ± 159.0 s; latency after stim+drug: 73.90 ± 34.84 s. p=0.0430, paired-t-test) (**Fig. S3C**).

**Figure 3:**
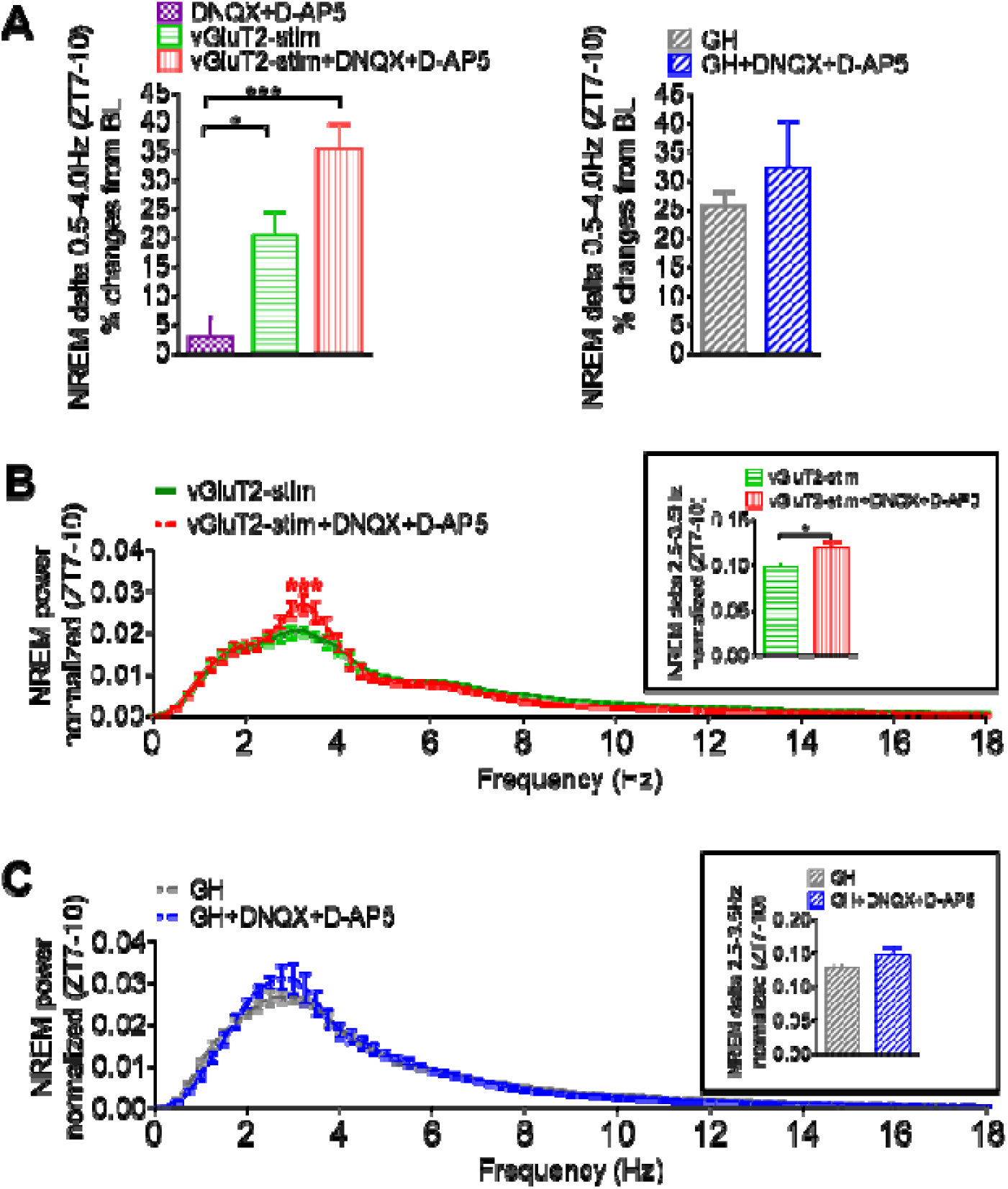
The increase in NREM sleep delta power after prolonged wakefulness induced by BF vGluT2-stim was comparable as that seen following gentle handling (GH) and was enhanced by local BF infusion of ionotropic glutamate receptor antagonists. **A**: Bar charts showing NREM delta power comparisons. vGluT2-stim and GH increased delta power (0.5-4 Hz) during NREM sleep compared to baseline (BL). Local applications of DNQX and D-AP5 did not affect NREM delta power (compared to baseline). *: p<0.05, ***: p<0.001, repeated-measures-ANOVA with Bonferroni’s comparison test. **B**: The increase of delta in the high frequency range (2.5-3.5 Hz) after vGluT2 stim was further elevated in the presence of drugs. Red asterisk: p<0.05 between vGlut2-stim and vGlut2-stim+DNQX+D-AP5. *: p<0.05, paired-t-test. C: Drugs did not affect NREM power spectra following GH. N=10 for vGluT2-stim, DNQX+D-AP5 and vGluT2-stim+DNQX+D-AP5 groups. N=7 for GH and GH+DNQX+D-AP5 groups.

### Whole brain cFos activation reveals differences in the activation of brain regions during SD induced by BF vGluT2-stim when compared to GH (**Figures 4**, 5)

The density of cFos immunostaining was significantly higher in 46 brain regions on the right hemisphere of the brain of mice in which the AAV injections and optostimulations were performed in right BF vGluT2 neurons (Table 1 green rows). Far fewer brain regions, most notably those involved in thalamocortical circuitry, showed statistically significant decreases in cFos density in the vGluT2-stim group compared to GH group (table 1 pink rows). A few other brain regions associated with sleep-wake modulation, such as the claustrum ^47^, VTA ^14^, and lateral hypothalamus ^48^ showed cFos activation in the vGluT2-stim group which was similar to that of GH group (table 1 gray rows). These differences are further depicted in heat maps showing average cFos density for each condition (**Fig 4A**). The subtracted density (Stim minus GH) of cFos in right coronal sections are plotted in **Fig 4B**. The magnified images of lateral septum (LS), bed nucleus of stria terminalis (BST), lateral habenula (LHb), striatum-like amygdalar nuclei (sAMY), basomedial amygdalar nucleus posterior (BMAp), tuberomammillary nucleus, ventral part (TMv), supramammillary nucleus (SUM), dorsal premammillary nucleus (PMd) and ventral premammillary nucleus (PMv) showed higher densities of cFos immunoreactivity with vGluT2-stim when compared to GH. These areas are closely associated with stress and avoidance, aversive and defensive behavior (**Fig 5A**). On the other hand, thalamocortical areas (somatosensory, SS and somatomotor, MO), parafascicular nucleus (PF), ventral group of dorsal thalamus (VENT), subparafascicular nucleus, parvicellular part (SPFp), dorsal part of the lateral geniculate complex, shell (LGd-sh), ventral part of the lateral geniculate complex (LGv), and posterior triangular thalamic nucleus (PoT) showed less cFos density with vGluT2 optostimulation than that with GH, suggesting reduced thalamocortical activation relative to GH (**Fig 5B**).We also observed that areas recognized for their involvement in sleep-wake behavior, such as ventral tegmental area (VTA), claustrum (CLA) and lateral hypothalamus (LH) show similar increases in cFos density when compared to undisturbed control (**Fig 5C**). The GFP protein mapping for locating the vGluT2 containing neuronal projection terminals, confirmed GFP fluorescence in the above listed brain regions (**Fig S4**).

**Figure 4:**
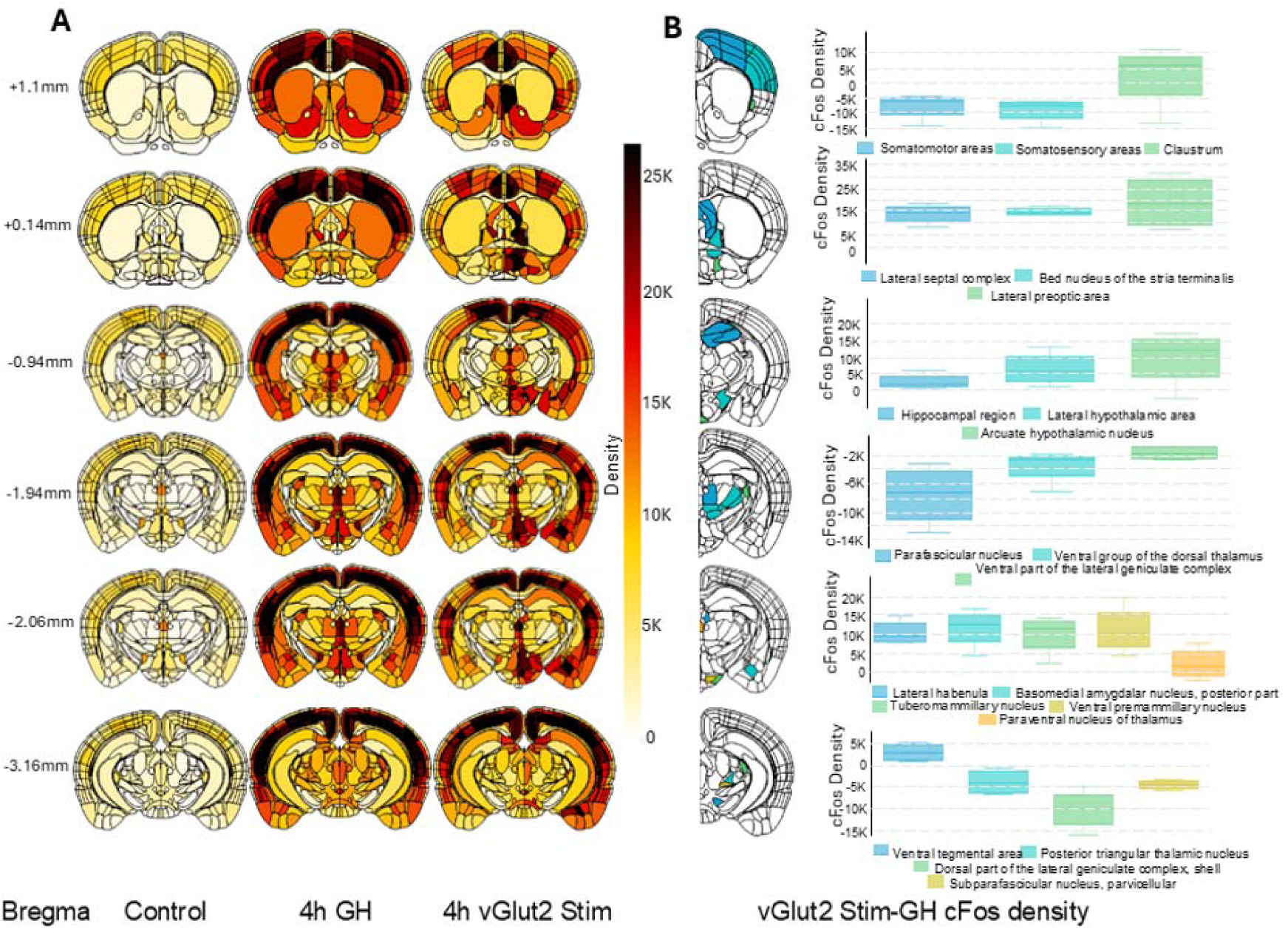
Comparison of cFos activation between GH and Optogenetic stimulation of BFvGluT2 neurons: **A**. Heat maps showing average density of cFos immunoreactivity in various brain regions of control (N=3), sleep deprivation with gentle handling (GH; N=4), and BF vGlut2 optostimulation (N=4) groups. **B**. Coronal sections of brain regions that showed either significantly increased, decreased or no change in cFos activity in BF vGluT2 stim when compared to GH group, the extent of the difference is plotted in the graph on the right side. All three groups were sacrificed at ZT7.

**Figure 5:**
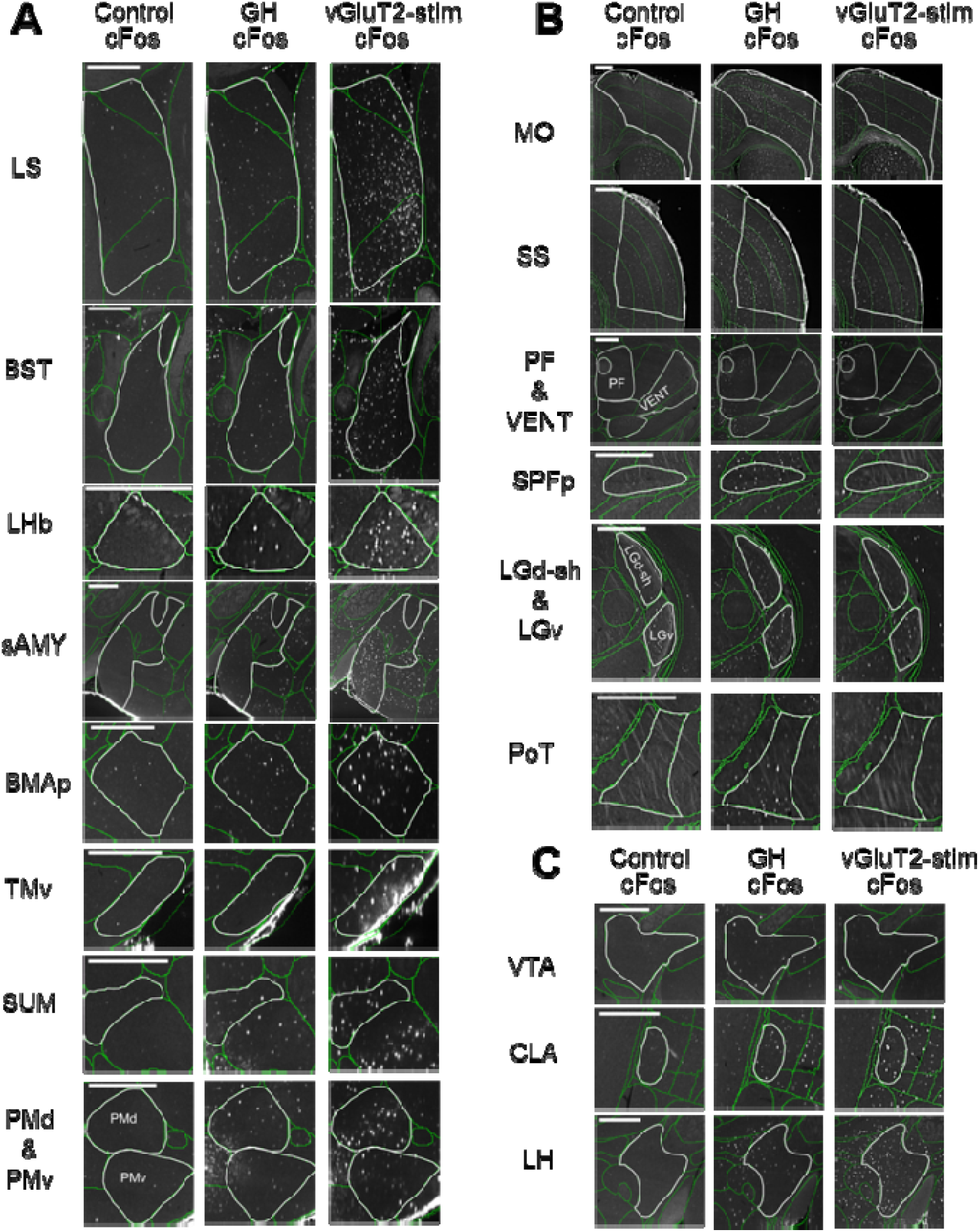
Immunohistochemical cFos protein labeling in brain regions that showed significant increases from the undisturbed sleeping control illustrated a different pattern of brain activity between gentle handling (GH) and optostimulation of BF vGluT2+ neurons (vGluT2-stim). **A.** Representative images showing regions with highest cFos positive cell density in vGluT2-stim group. **B.** Representative images showing regions with highest cFos positive cell density in GH group. **C.** Some brain regions implicated both in sleep-wake and motivated behavior showed elevated cFos positive cells compared to the control but did not show statistically significant difference in cFos between GH and vGlut2-stim groups. Both groups showed higher cFos immunolabeling compared to controls. Scale bar 500 µm. LS: Lateral septum. BST: Bed nuclei of the stria terminalis. LHb: Lateral habenula. sAMY: Striatum-like amygdalar nuclei. BMAp: Basomedial amygdalar nucleus, posterior. TMv: Tuberomammillary nucleus, ventral part. SUM: Supramammillary nucleus. PMd: Dorsal premammillary nucleus. PMv: Ventral premammillary nucleus. MO: Somatomotor areas. SS: Somatosensory areas. PF: Parafascicular nucleus. VENT: Ventral group of the dorsal thalamus. SPFp: Subparafascicular nucleus, parvicellular part. LGd-sh: Dorsal part of the lateral geniculate complex, shell. LGv: Ventral part of the lateral geniculate complex. PoT: Posterior triangular thalamic nucleus. VTA:Ventral tegmental area. CLA: Claustrum. LH: Lateral hypothalamic area.

**Table 1:**
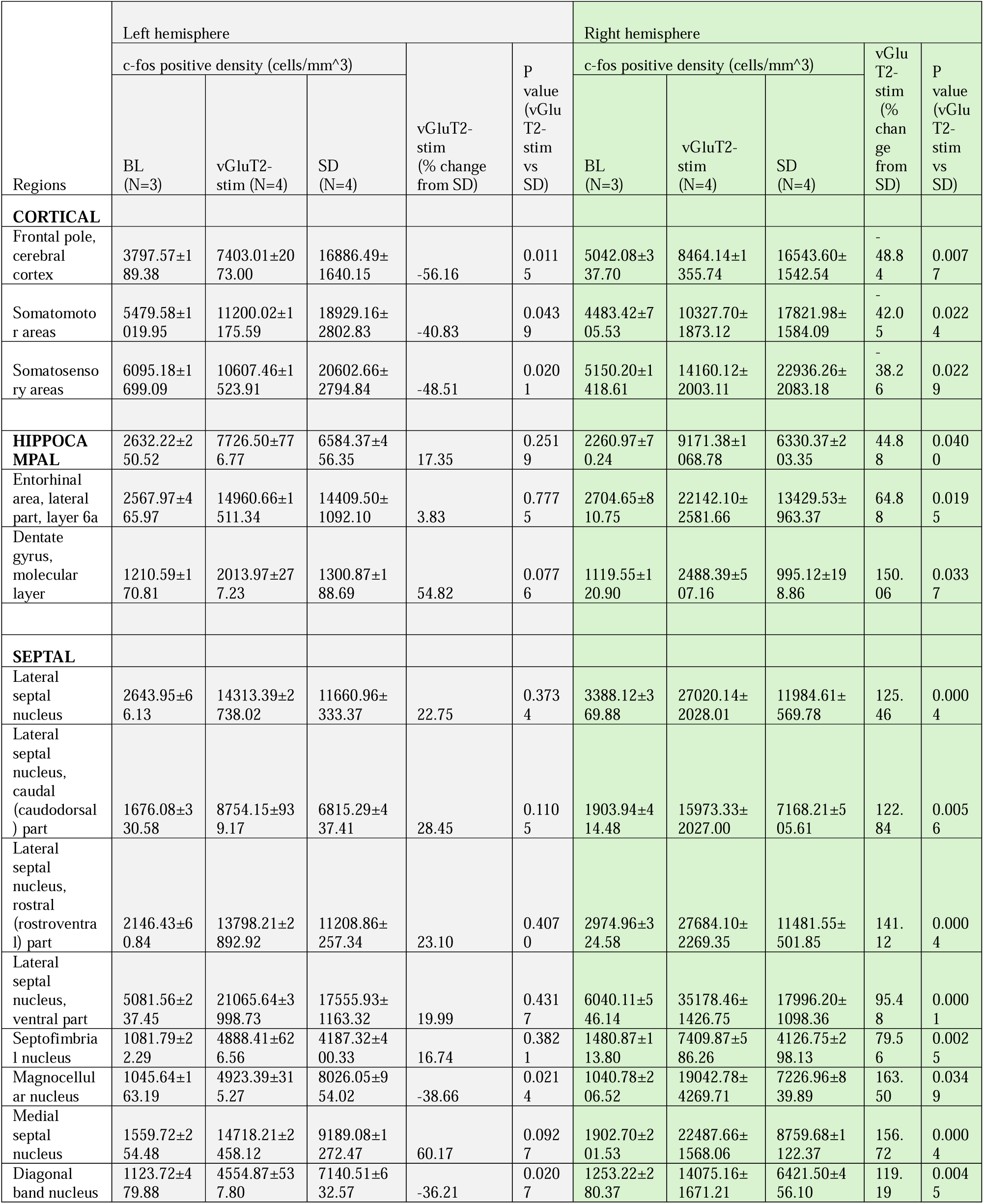
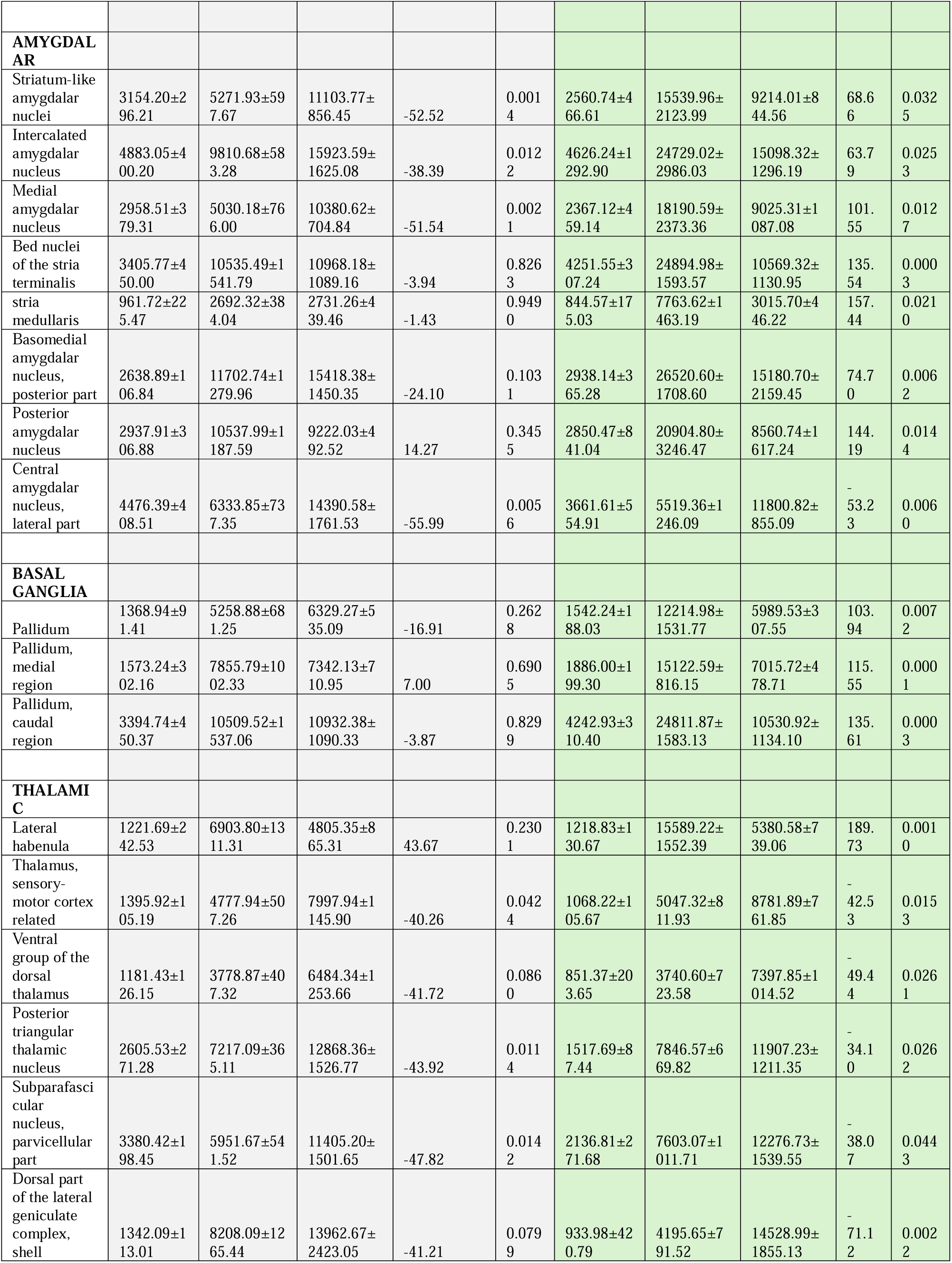
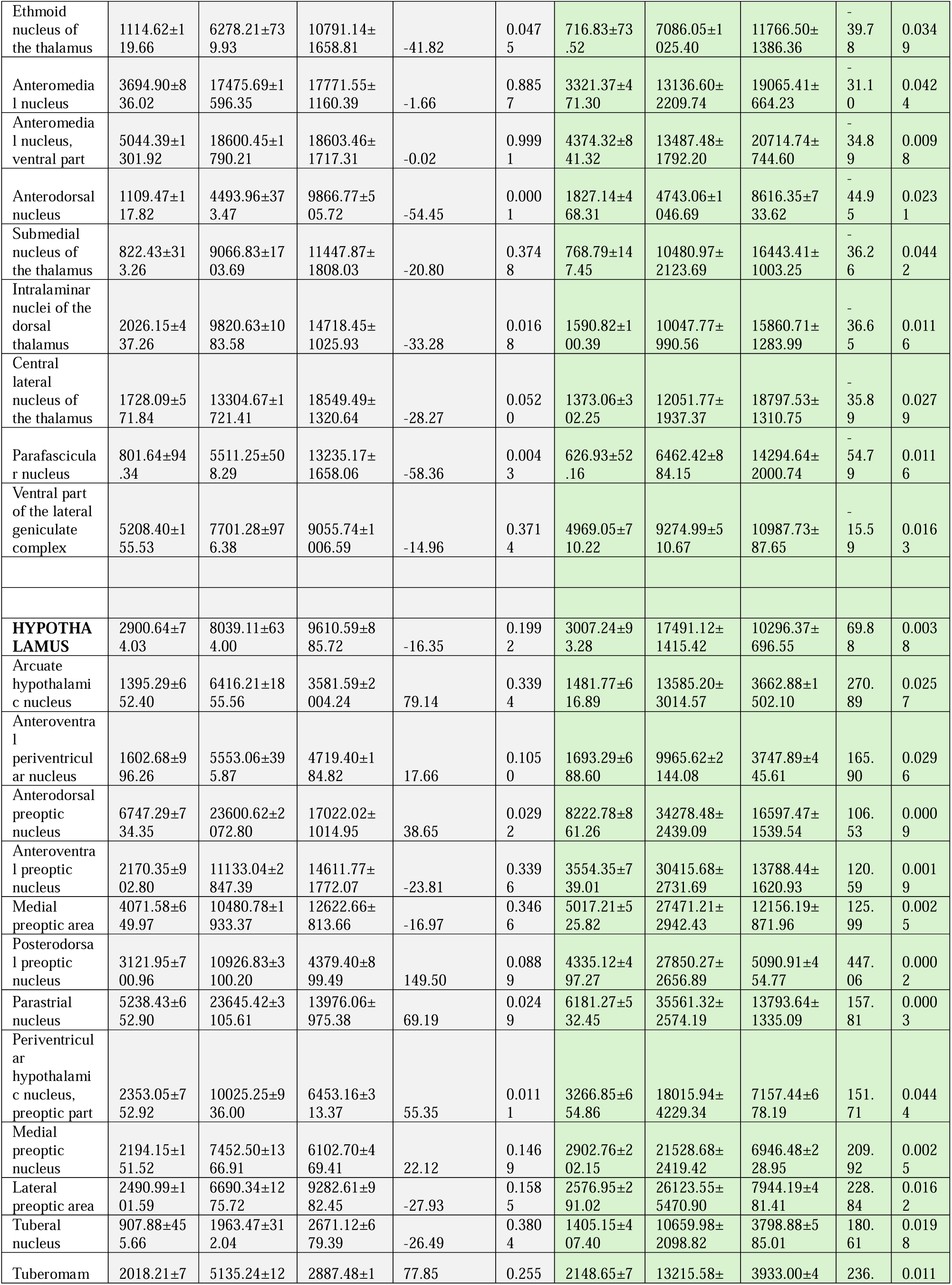
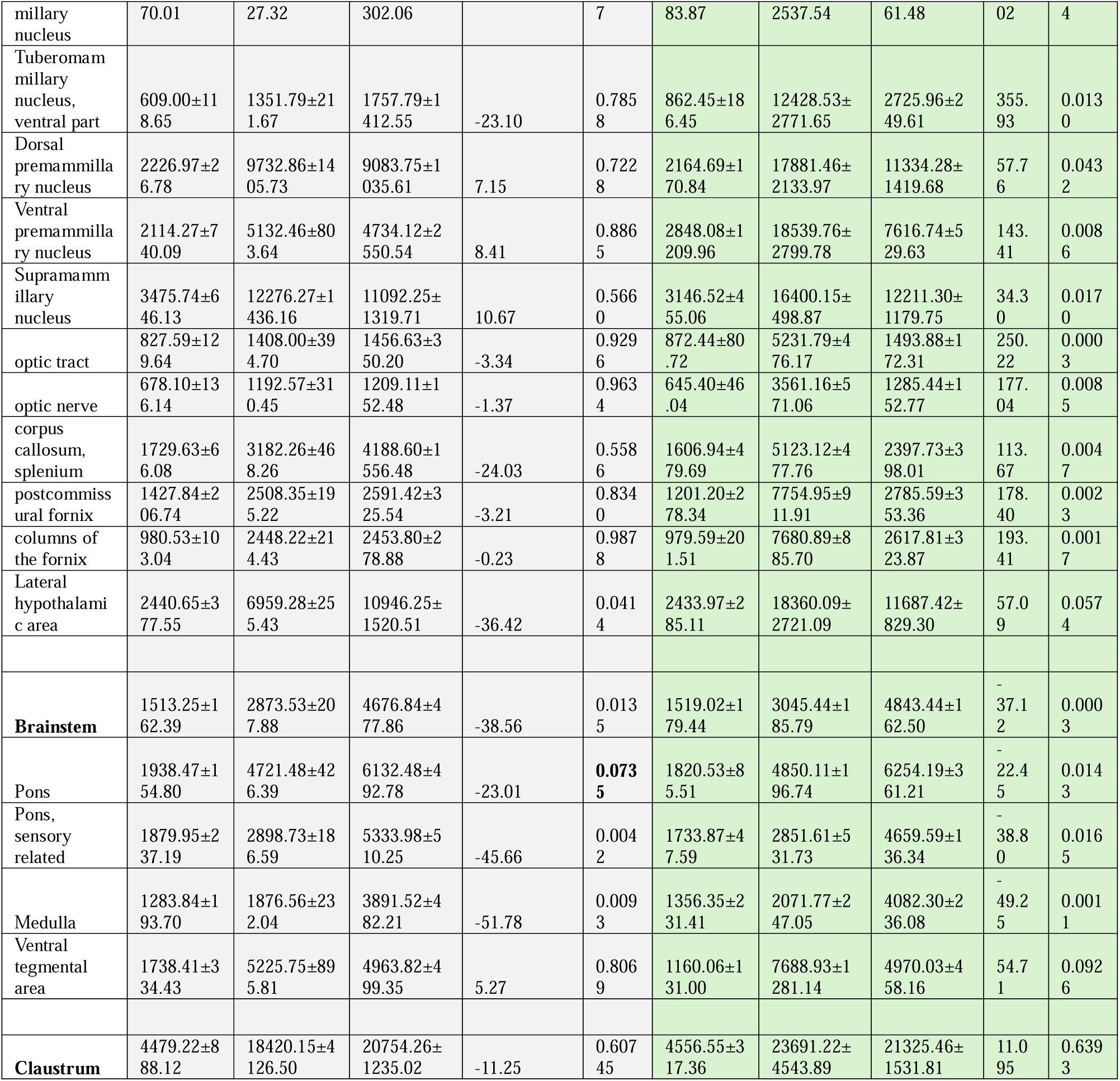
List of brain regions with c-fos positive cell density and comparison between vGluT2-stim and SD.

## Discussion

Considerable evidence supports a role for the BF in promoting wakefulness and the HSR. Selective optogenetic or chemogenetic activation of all three major BF neurotransmitter subtypes, GABAergic, cholinergic, and glutamatergic neurons, can enhance the amount of time spent in wakefulness ^23,34,35,49–54^. Activation of BF cholinergic neurons can enhance the amount of wakefulness ^35,52^ through excitation of other BF subtypes ^35,55^ so also BF GABAergic/parvalbumin (PV)-expressing cells, ^50^, BF GABAergic neurons expressing the transcription factor neuronal Per-Arnt-Stim (Npas1) ^54^ and vGluT2 neurons ^34^. Cholinergic neurons are also associated with the HSR and with the extracellular release of homeostatic sleep-promoting substances such as adenosine and nitric oxide ^56,57^. More recently, BF vGluT2 neurons have also been implicated in short-term (seconds or minutes long), activity-dependent local increases in extracellular adenosine and in homeostatic processes ^58^. However, while selective lesion of cholinergic neurons attenuates sleep deprivation-induced increases in adenosine and EEG responses to sleep deprivation ^56^, such loss-of-function tests were not performed in sleep deprived mice with BF glutamatergic neuronal lesion ^58^. Thus, the role of BF vGluT2 neurons in the HSR remains an open question. A direct comparison of the HSR produced by activation of BF vGluT2 neurons and that produced by a widespread method for sleep deprivation in rodents, GH, has not previously been performed. Thus, here we have compared the wake and recovery sleep EEG parameters for 4h of GH-and vGluT2-stim-induced wakefulness. Our results show that 4h of vGluT2 neuronal stimulation during the early light period led to wakefulness comparable to that observed with time-matched 4h of GH. However, responses to locally administered glutamate receptor antagonists during vGluT2-stimulation, and differences in certain EEG parameters, suggested a predominance of local glutamate action on neighboring wake-promoting neurons and associated circuitry which differed from that of GH-induced wakefulness. Differences were also noted in the EEG biomarkers for sleep homeostasis during waking and the ensuing sleep, suggesting a weaker homeostatic response in the vGluT2-stimulation group compared to the GH group. Comparison of brain c-Fos activity revealed stronger neuronal activation in brain regions involved in avoidance, aversive and defensive behavior with vGluT2 stimulation, consistent with recent studies of the function of these neurons ^22–26^. In contrast, GH showed higher activation in regions associated with thalamocortical arousal circuitry.

Within BF, vGluT2 neurons send excitatory projections to intra-BF GABAergic (parvalbumin- and somatostatin-expressing) as well as cholinergic neurons ^34^. BF glutamatergic neurons promote arousal though local excitation of other BF wake-promoting subpopulations, including parvalbumin-expressing GABAergic, cholinergic ^34^, and likely also Npas-1 positive neurons ^54^, as well as through direct projections to other arousal-related brain regions ^23^. The differences in the effects of locally applied glutamate receptor antagonists on the wakefulness elicited by vGluT2-stim indicated predominance of selective BF glutamatergic neuronal circuitry-mediated effects on arousal, when compared with GH (Fig 2B1 and 2; & Fig S1). The vGluT2-stim-induced wakefulness, although not significantly different from that of GH, showed a distinct pattern of gradual hourly decrease, a pattern that might reflect the excitation of neighboring cholinergic ^34^ involved in the release of homeostatic sleep promoting factors such as adenosine ^39,56,58^. While a similar increase in adenosine is also observed during GH ^59^, the effect on wakefulness may be partially masked by activation of other wake-promoting systems. The local antagonism of glutamate receptors enhances the decline in wakefulness significantly, suggesting the local action of glutamate within BF in promoting wakefulness.

The spectral analysis of the EEG during wakefulness showed a pronounced and significant increase in delta activity ranging from 0.5-3Hz in the vGluT2-stim group. The delta increases in GH group was restricted to 1-1.5Hz. Increased wake delta has been observed previously in rodent paradigms which disrupt sleep, such as sleep interruption ^60^. Increased delta is also associated with aspects of wakefulness such as avoidance behavior in response to pain or anxiety. Individuals with higher pain sensation scores showed a greater increase in wake delta, suggesting that avoidance behavior manifests through increased delta power needed to inhibit painful sensory afferences ^61^. An increase in wake delta has also been reported in patients with anxiety disorders ^62^. It is suggested to be due to the breakdown in top-down regulation, where the subcortical emotional centers (like amygdala) override or suppress conscious prefrontal cortical activity leading to avoidance behavior ^63^. Our results support these ideas, given higher activation in avoidance behavior-related brain areas with vGluT2-stim, as also indicated by cFos immunohistochemistry results discussed below.

Wake EEG theta (5-9 Hz) activity increased significantly for both the vGluT2-stim and GH groups. Cortical theta activity depends upon input from BF cholinergic neurons ^64,65^. Interestingly, the low theta (5-7 Hz) frequency associated with homeostatic sleep drive ^38,39^ showed no change in the vGlut2-stim group but showed a trend-level increase in the GH group, suggesting an increased homeostatic drive in the latter but not the former. Significant increases in the high theta (7-9 Hz) range, associated with wake-associated movements, were blocked by local administration of glutamate receptor antagonists. Targeted microinjections of glutamate agonists into BF increase cortical theta activity attributed to its action on neighboring cholinergic neurons’ bursting discharge ^66–68^. Our results confirm the effect of BF vGluT2-stim on cortical theta activity is due to a local effect on neighboring BF neurons since it was blocked by local administration of glutamate receptor antagonists.

The NREM sleep amount failed to show any increases following 4 h of vGluT2-stim. Moreover, the increase in NREM delta (0.5-4 Hz) activity in the ensuing sleep was far less pronounced in the vGluT2-stimulation (20.09% increase), than the GH group (53.65%) when compared to their respective baselines. Taken together, absence of any increases in homeostatic wake low theta (5-7Hz), and recovery NREM sleep, and a moderate increase in NREM delta suggest a weaker homeostatic response after BF vGluT2-stimulation-induced arousal compared to GH-induced arousal.

Activation of different arousal pathways have been shown to elicit different homeostatic sleep responses ^12–18^. Examining the whole brain cFos activation pattern suggests that vGluT2-stim led to notably higher activation of brain areas associated with avoidance, aversive and defensive behavior when compared to GH, as expected given the projections of these neurons. Our group and others have shown that BF vGluT2+ neurons project to the frontal cortex, lateral hypothalamus, amygdala, lateral habenula, lateral hypothalamus, VTA and other brain areas involved in the control of arousal, reward/aversion and defensive behavior ^23,30,69–72^. Recently, Zou et al., (2024)^73^ performed vGluT2 optogenetic stimulation in conjunction with functional magnetic resonance imaging (fMRI) in awake mice, and showed that the strongest and most widespread BOLD responses occurred in the areas previously shown to be involved in wakefulness and avoidance/defensive behavior ^23,25,34,58,74^. Our results on whole-brain cFos activation provide anatomical evidence in support of these reports, showing that higher cFos activation occurs in similar brain areas when compared to the GH group. In addition, we observed that the LS, BST, LHb, aAMY, BMAp, TMy, SUM, PMd, and PMv, all show higher density of cFos immunolabeling compared to GH. Collectively, these regions respond to aversive stimuli leading to avoidance behavior ^75,76^. For example, LS is recognized as a critical node in the regulation of emotional behavior in response to aversive experiences, leading to avoidance ^77^. The SUM recently emerged as a potential new component of the behavioral avoidance system ^78,79^. Lesions of SUM are associated with reduced avoidance in the elevated T-maze. Basal forebrain glutamatergic neurons project prominently to LHb ^23,80^. The LHb is a key brain center for anti-reward-based decision-making, negative reinforcement, and avoidance ^81,82^. Cross-species behavioral research (humans with calcified BLA/BMA and chemogenetically silenced BLA/BMA) revealed that under imminent threat, the BLA/BMA is essential for rapid avoidance behavior ^83^.

We also observed that brain regions constituting the thalamocortical circuitry and prominently involved in sleep-wake regulation showed higher cFos immunolabeling in GH group, compared to the vGluT2-stimulation group, a possible mechanistic observation related to the higher NREM delta activity we observed in the GH group during the recovery period. Interestingly, the cFos activity was equally elevated in areas known to be involved in arousal as well as avoidance behavior, such as VTA ^14,84,85^, claustrum ^86,87^, and lateral hypothalamus ^88,89^. Thus, following the pattern of cFos activation, our results set the stage for identifying brain regions specifically involved in wakefulness and/or avoidance/defensive behavior. The higher activation of brain regions involved in avoidance behavior in BF vGluT2-stimulation-induced arousal showed an attenuated homeostatic response, likely a necessity, preventing excessive sleepiness during the defensive/avoidance state.

Our results indicate that BF-vGluT2 neurons induce arousal by activating the avoidance/defensive behavior circuitry with an associated weak homeostatic sleep response. These behaviors are often used to model human illnesses; for example, insomnia is often conceptualized as a disorder of hyperarousal ^19^ and anxiety can be viewed as an excessive avoidance response to potential or actual threats, thus highlighting a potential target for treatment of human disorders that elicit avoidance behavior. Many psychiatric disorders involve excessive avoidant or defensive behavior, such as avoidance in anxiety and trauma disorders or defensive rituals in obsessive-compulsive disorders. Anxiety and trauma also exacerbate arousal and weaken homeostatic response for sleep rebound. From an evolutionary perspective these changes in the HSR may serve as an adaptation which would maintain enhanced vigilance in the face of an ongoing or perceived threat.

## STAR METHODS

### Resource Availability Lead Contact

Further information and requests for resources and reagents should be directed to and will be fulfilled by the Lead Contact, Radhika Basheer.

### Materials Availability

This study did not generate new unique reagents.

### Data and Code Availability

The datasets supporting the current study have not been deposited in any public repository but are available from the lead author upon request. This study did not generate new code.

### Experimental Model and Subject Details Animals

Adult (3-6 months old) male vGluT2-cre mice (congenic strain 028863, Jackson labs, Bar Harbor, ME, USA) were used for the *in vivo* optogenetic, opto-dialysis, sleep recording and c-Fos staining experiments. In addition, Adult (3-6 months old) male C57Bl6 mice (# 000664, Jackson Labs) were used for the baseline sleep recordings, sleep deprivation and pharmacological interventions by reverse microdialysis. Mice were housed under constant temperature and a 12h:12h light/dark cycle (Lights on, ZT0=7:00AM; Lights Off, ZT12=7:00PM), with food and water available *ad libitum*. The Institutional Animal Care and Use (IACUC) and Biosafety Committees of VA Boston Healthcare System approved the animal experiments described herein. All experiments were performed in accordance with the National Institute of Health Guide for the Care and Use of Laboratory Animals (NIH Publications No. 80–23) and conformed to US Veterans Administration, Harvard University, and the US National Institutes of Health guidelines. All efforts were made to minimize the number of animals and their suffering.

### Stereotaxic surgery for AAV5-DIO-ChR2-EYFP injection and implantation of opto-dialysis/or reverse microdialysis probe or optical fiber, and EEG/EMG headmounts

All mice (C57Bl6 and vGluT2-Cre mice) underwent stereotaxic surgery. Mice were deeply anesthetized with isoflurane (induction: 3-5%; maintenance: 1.5%– 2.5%) and placed in a stereotaxic device (Kopf Instruments). Both C57Bl6 and vGluT2-cre mice were implanted with EEG/EMG head mounts Catalogue #8201, Pinnacle Technology Lawrence, KS, USA) in the same animals to monitor the sleep-wake states. The EEG recording electrodes were implanted into the skull above the frontal cortex (coordinates: AP: 2.2mm, ML ±0.5mm). The reference electrode was placed above the cerebellum. The ground electrode was placed above the temporal cortex. EMG electrodes were implanted in the dorsal neck musculature to monitor the muscle tone. The C57Bl6 mice were bilaterally implanted with microdialysis CMA 7 guide cannulae (Harvard Apparatus) targeting the BF regions (AP: 0; ML: -1.5mm; DV: -5.6). After 1 week of recovery from the surgery, these mice were used for BL sleep recordings and sleep deprivation with GH with and without reverse microdialysis of the ACSF/cocktail of ionotropic glutamate receptor antagonists described below. The vGluT2-Cre mice received injections of 0.7 μl AAV5-DIO-ChR2-EYFP into the BF region of the right hemisphere, followed by implantation of our opto-dialysis probe (a combination of optical fiber and microdialysis probe) ^35^ or optical fiber (Doric Lenses, Inc, Quebec City, Canada) targeting the same BF region. For C57Bl6 mice the experiments were performed one week after the recovery from the surgery. Whereas, for vGluT2-Cre mice the experiments were done one month after the surgery in order to allow the viral transduction and expression in vGluT2 neurons. Mice were tethered to the sleep recording device in the recording room for at least two days before the experiments and were allowed 2 days of recovery between experiments.

### Optical stimulation

A laser light supply device (MBL-III-473 nm-50mW,OptoEngine) linked to the optic fiber was used to provide 473nm-wavelength light. The following stimulation paradigm was used in this study: 20Hz, 10ms pulse, 20mW/cm^2^, 5 s per min.

### In vivo drug delivery by reverse microdialysis

A CMA 7 1 mm microdialysis probe (Harvard Apparatus, Cambridge, MA, USA) was inserted into the guide cannula at least 12 h before the start of infusion experiments. The inlet of the microdialysis probe was linked to a CMA 400 Syringe pump (Harvard Apparatus) via microdialysis tubing which allowed the drugs to be infused continuously at a speed of 1 µl/min. Artificial cerebrospinal fluid (ACSF; Harvard Apparatus) infusion and non-infusion baseline recordings served as controls. A random and counter-balanced sequence of different testing conditions (drug infusion, ACSF infusion or baseline) was applied in the experimental design to avoid order effects. The ionotropic glutamatergic receptor antagonists 200 μM DNQX (Cat. # 505026, Millipore Sigma, Burlington MA) and 500 μM D-AP5 9 Cat # A8054, Millipore Sigma, Burlington MA) were used in this study.

### Sleep deprivation (SD) by gentle handling (GH)

Sleep deprivation was performed using the gentle handling protocol as previously reported by us and others ^56,59,90–92^. In brief, mice were kept awake by introducing novel objects into the cages, replacing part of bedding with fresh bedding and gentle touching with soft tissue or gloves. SD was performed for 4 hours (ZT3-7). The EEG/EMG was continuously recorded to ensure the effectiveness of SD. Mice were allowed 2 d to recover before the next set of experiments.

### EEG/EMG recording scoring and analysis

EEG/EMG polygraphic data were recorded with a Pinnacle 3-channel 8200 system (Pinnacle Technology). The signals were captured with Sirenia Acquisition program (Pinnacle Technology) at a sampling rate of 1 kHz. The frontal EEG recording of the right hemisphere was used for sleep-state analysis and power spectral analysis. Sleep-wake states were scored manually offline in 4-s epochs using Sirenia software (Pinnacle Technology). EEG epochs without movement artifacts were used for power spectral analysis using Sirenia software. Power spectral analysis was performed with Sirenia Sleep software for each state-dependent frequency band and then normalized to the value of total power from 0-18 Hz for all states for the respective time window (wake, NREM, and REM). For example, wake power from ZT 3-7 was normalized to the value of total power from 0-18 Hz from ZT3-7. NREM power from ZT7-10 was normalized to the value of total power from 0-18 Hz from ZT7-10.

### Neuronal activity examination with whole-brain c-Fos mapping

vGluT2-cre mice (2-3 month-old) were randomly separated into three groups. Based on the following experimental design, each mouse received one surgery for inserts/implantations following the stereotaxic surgery method described above. The control group received the EEG/EMG headmount implantation (N=3). The opto-stimulation group was injected with AAV5-DIO-ChR2-EYFP into the right BF followed by insertion of the optical fiber targeting BF as well as implantation of EEG/EMG headmount (N=4). The sleep deprivation (SD) group received the implantation of EEG/EMG headmount (N=4). During the experiment day, the control group was allowed to sleep/wake spontaneously. The opto-stimulation group received 4h optical stimulation (ZT3-7). The SD group was sleep deprived for 4 hours (ZT3-7). EEG/EMG signals were recorded from all animals during ZT3-7 to monitor the sleep-wake states. At ZT7, the mice were transcardially perfused with PBS containing 10U/ml heparin, followed by 10% formalin overnight and transferred to PBS with 0.02% sodium azide. Tissues were processed by LifeCanvas Technologies (Cambridge, MA) for c-Fos staining, imaging and analysis. In brief, brains were processed following the SHIELD protocol as previously published ^93,94^. The samples were cleared for 7 days with Clear+delipidation buffer, followed by staining using the SmartBatch+ system with c-Fos antibody (Abcam ab214672, secondary antibody as SeTau-647 which showed red fluorescence). The labeled samples were incubated in EasyIndex (LifeCanvas Technologies) for regractive index matching (n=1.52) and the images were captured with light sheet microscope SamrtSPIM and post-processed with SmartAnalytics for cell counts, densities, heat maps and graphical outputs. Because of technical limitations raised by resolution limitations and distinct subcellular expression patterns of GFP (cytosolic or projection terminals) and FOS (nuclear) signals in a three-dimensional space, the direct colocalization between FOS+ and GFP+ cells was not analyzed.

### Statistical analysis

Data were presented as the mean ± SEM. ANOVA test followed by a Bonferroni correction was used for comparisons among multiple groups. Repeated measures ANOVA followed by Bonferroni’s correction was used for comparisons among multiple sets of dependent data. T-test was used to compare differences between two groups. Paired-t-test was used to compare difference between two sets of dependent data. GraphPad Prism software (GraphPad Software, Inc. La Jolla, CA, USA) was used for statistical tests. Differences were considered significant at *p* < 0.05.

## Key Resource Table

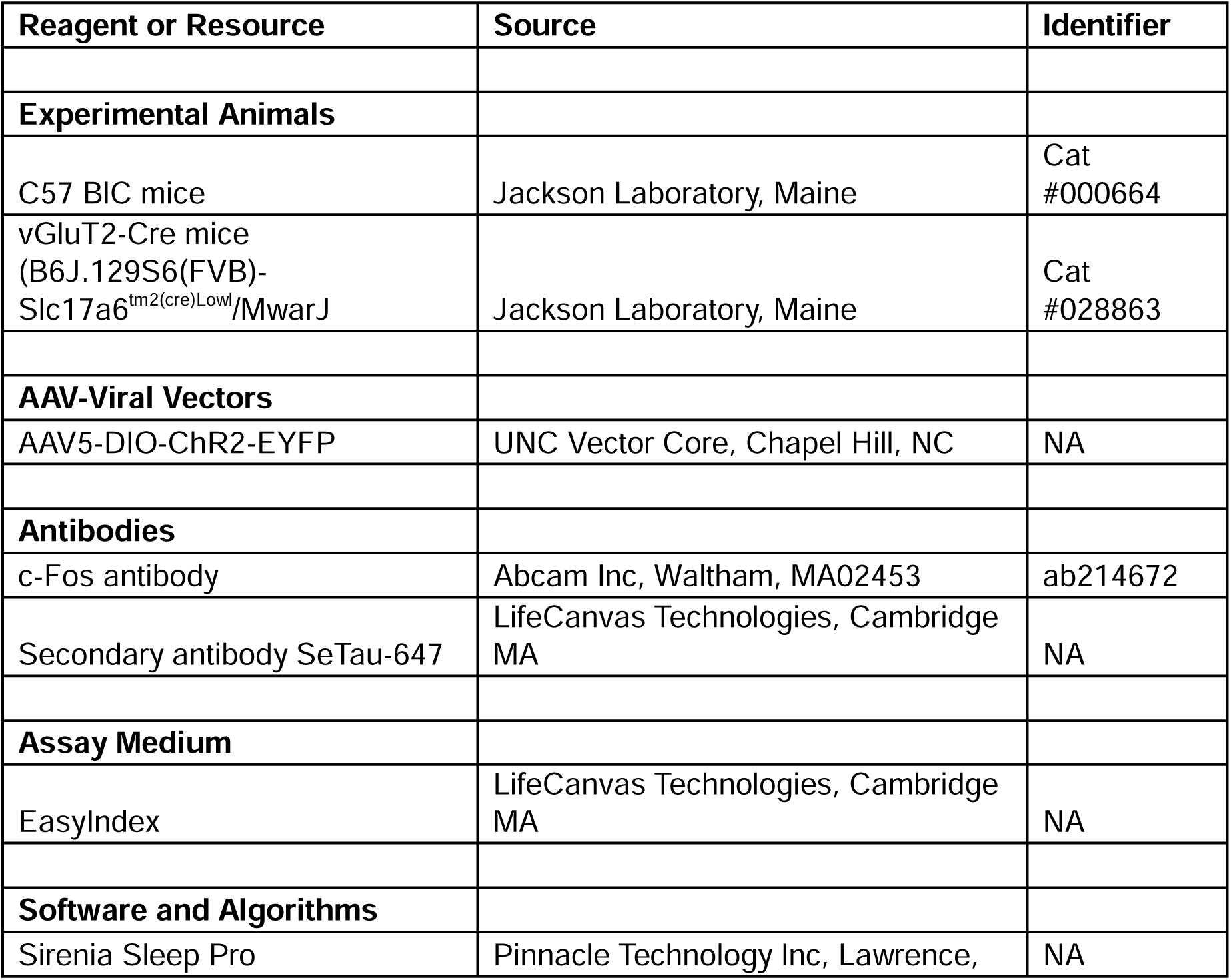
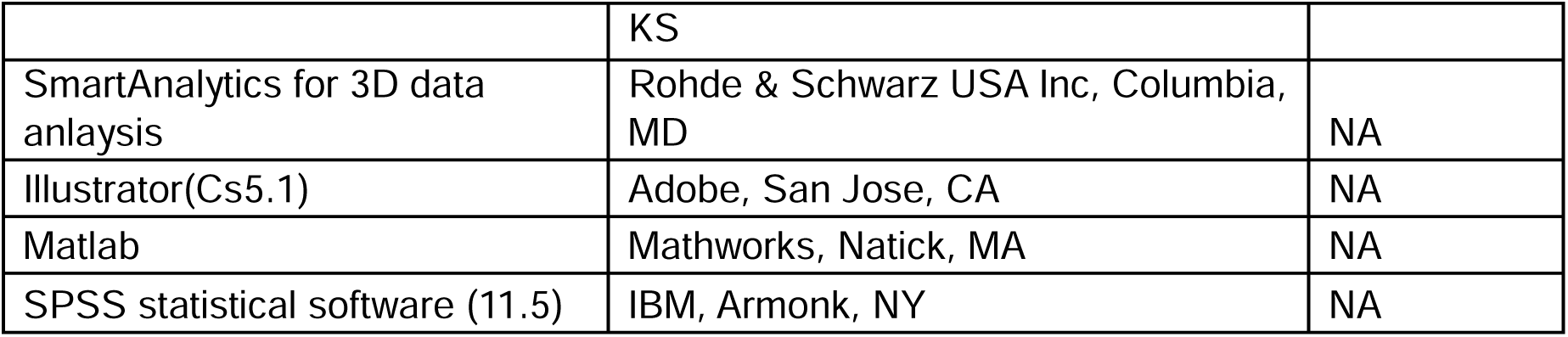

## Supporting information

NIHPP102026

## ACKNOWLEDGMENT

This work was supported by National Institutes of Health grant R01-NS119227 (RB), VA Biomedical Laboratory Research and Development Service Merit Awards, I01 BX001404 & I01 BX006105 (RB), I01 BX004673 (REB), CDA IK2 BX004905 (DSU). JTM received partial salary compensation and funding from Merck MISP (Merck Investigator Sponsored Programs) but has no conflict of interest with this work. RB, JTM and DSU and REB are Research Health Scientists at VA Boston Healthcare System, West Roxbury, MA. All work was conducted at VA Boston Healthcare System. The contents of this work do not represent the views of the U.S. Department of Veterans Affairs or the United States Government.

## AUTHOR CONTRIBUTIONS

C.Y., R.E.B., & R.B. conceived and designed the experiments. C.Y., J.T.M., & V.L. performed the experiments. C.Y., J.T.M., D.S.U., & R.B. analyzed the data. C.Y., J.T.M., R.E.B. & R.B. drafted and revised the manuscript. Other authors provided comments on the manuscript.

