## Supplementary material for "Sustained Activation of a Defensive Behavior-Related Circuit Elicits a distinct Homeostatic Sleep Response": NIHPP102026

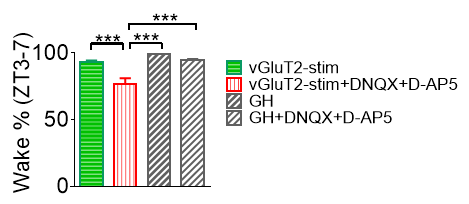

**Supplemental figure 1:** Local blockade of ionotropic glutamate receptors in BF reduced the increase in wakefulness induced by optostimulation of BF vGluT2+ neurons (vGluT2-stim), but not gentle handling (GH). ***: p<0.001, ANOVA with Bonferroni’s comparison test.

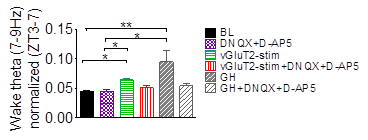

**Supplemental figure 2**: Optostimulation of BF vGluT2+ neurons (vGluT2-stim) and sleep deprivation by gentle handling (GH) increased power of high theta activity (7-9 Hz) in the wake state compared to baseline (BL). Local blockade of ionotropic glutamate receptors in BF showed a trend decrease in theta activity induced by vGluT2-stim or GH. *: p<0.05, **: p<0.01, ANOVA with Bonferroni’s comparison test .

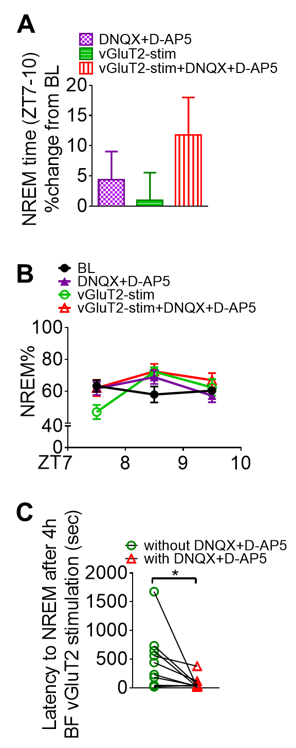

**Supplemental figure 3:** Blockade of BF ionotropic glutamate receptors does not alter the percentage of time spent in NREM sleep during the recovery period but shortens the latency to NREM after BF vGluT2 -stim-induced prolonged wakefulness. A. Summary of NREM time during the recovery period ZT7-10. vGluT2-stim did not induce an elevation of NREM sleep during the recovery period. In the presence of drugs (DNQX+D-AP5), NREM sleep after vGluT2-stim showed an upward trend. There was no statistically significant difference compared with baseline (BL).N=10. **B**: Hourly NREM time % during the recovery period. During the first recovery hour, NREM % after vGluT2-stim was low, compared to other groups. **C**: Latency to NREM after 4h BF vGluT2-stim was significantly decreased in the presence of ionotropic glutamate receptor antagonists applied to BF. *: p<0.05. paired-t-test.

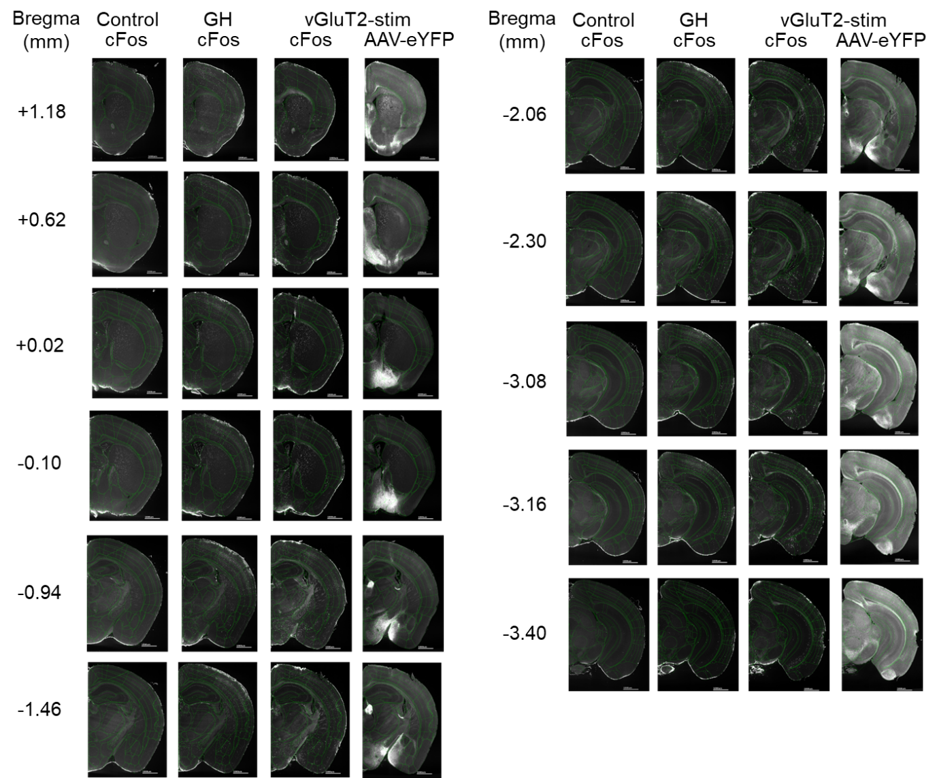

Supplemental figure 4: Coronal sections showing neuronal activity (cFos expression) in similar location to the BF vGluT2 neurons projection areas (AAV-DIO-CHR2-eYFP). Baseline: Control; GH: sleep deprivation by gentle handling; vGluT2-stim: BF vGluT2 optostimulation. AAV-eYFP: GFP fluorescence (bright) induced by AAV5-DIO-ChR2-eYFP injection in the BF of vGluT2-cre mice.
